# Non-autophagy role of Atg5 and NBR1 in unconventional secretion of IL-12 prevents gut dysbiosis and inflammation

**DOI:** 10.1101/2020.12.07.414227

**Authors:** Seth D. Merkley, Samuel M. Goodfellow, Yan Guo, Zoe E.R. Wilton, Janie R. Byrum, Kurt C. Schwalm, Darrell L. Dinwiddie, Rama R. Gullapalli, Vojo Deretic, Anthony Jimenez Hernandez, Steven B. Bradfute, Julie G. In, Eliseo F. Castillo

**Author notes:** Authors contributed equally to this study (Seth D. Merkley, Samuel M. Goodfellow). Correspondence should be addressed to (E.F.C.).

## Abstract

Intestinal myeloid cells play a critical role in balancing intestinal homeostasis and inflammation. Here, we report that expression of the autophagy related 5 (Atg5) protein in myeloid cells prevents dysbiosis and excessive intestinal inflammation by limiting IL-12 production. Mice with a selective genetic deletion of *Atg5* in myeloid cells (Atg5ΔMye) showed signs of dysbiosis prior to colitis and exhibited severe intestinal inflammation upon colitis induction that was characterized by increased IFNγ production. This increase in IFNγ was due to excess IL-12 secretion from *Atg5*-deficient myeloid cells. Atg5 functions to limit IL-12 secretion through modulation of late endosome (LE) acidity. Additionally, the autophagy cargo receptor NBR1, which accumulates in Atg5-deficient cells, played a role by delivering IL-12 to LE. Restoration of the intestinal microbiota and alleviation of intestinal inflammation was achieved by genetic deletion of IL-12 in Atg5ΔMye mice. In summary, Atg5 expression in intestinal myeloid cells acts as an anti-inflammatory brake to regulate IL-12 thus preventing dysbiosis and uncontrolled IFNγ-driven intestinal inflammation.

## INTRODUCTION

Immune system dysregulation, intestinal barrier defects and dysbiosis are believed to be driven in part by genetics according to recent genome-wide association studies (GWAS) (Schirmer, Garner et al., 2019). Many of the identified genes are involved in innate cell bacterial recognition and processing and appear to contribute to the pathogenesis observed in inflammatory bowel disease (IBD) such as *NOD2* and *ATG16L1* which are linked to autophagy (Schirmer et al., 2019). Autophagy is one such process associated with IBD susceptibility (Brest, Lapaquette et al., 2011, Hampe, Franke et al., 2007, Henckaerts, Cleynen et al., 2011, Lahiri, Hedl et al., 2015, Lassen & Xavier, 2018, Levine & Kroemer, 2019, McCarroll, Huett et al., 2008, Rioux, Xavier et al., 2007, Roberts, Gearry et al., 2007, Yamazaki, Onouchi et al., 2007). Autophagy is a conserved catabolic process that degrades protein aggregates, damaged organelles and numerous pathogens (Galluzzi, Baehrecke et al., 2017). Autophagy has proven critical for intestinal homeostasis. Defects in the autophagic pathway, specifically in intestinal epithelial cell lineages, results in increased intestinal permeability and Paneth and Goblet cell (GC) dysfunction (Cadwell, Liu et al., 2008, Cadwell, Patel et al., 2009, Matsuzawa-Ishimoto, Shono et al., 2017, Nighot, Hu et al., 2015, Patel, Miyoshi et al., 2013, Wong, Ganapathy et al., 2019). More recently, an IBD risk loci associated with autophagy was found to disrupt the microbiota albeit it is unclear what cell-type mediates this effect (Lavoie, Conway et al., 2019). Nevertheless, there is mounting evidence that autophagic genes are critical for intestinal homeostasis and defects in the autophagic process can lead to increased susceptibility to intestinal pathogens and overall enhanced IBD susceptibility (Bel, Pendse et al., 2017, Benjamin, Sumpter et al., 2013, Burger, Araujo et al., 2018, Lassen & Xavier, 2018).

Many of these IBD risk genes and pathways including autophagy are highly relevant to myeloid cell function in addition to that of epithelial cells (Chauhan, Mandell et al., 2015, Homer, Richmond et al., 2010, Singh, Davis et al., 2006, Zhang, Zheng et al., 2017). However, the specific role for autophagy and autophagy genes in myeloid cells in maintaining the balance between intestinal homeostasis and inflammation has yet to be fully explored. The prevailing hypothesis linking autophagy to IBD is through the IL-17 signaling pathway. IL-1, IL-17 and IL-23, all involved in IL-17-mediated inflammation, are upregulated in IBD (Angelidou, Chrysanthopoulou et al., 2018, Coccia, Harrison et al., 2012, Dragasevic, Stankovic et al., 2018, Jiang, Su et al., 2014, Kanai, Mikami et al., 2012, Liu, Yadav et al., 2011, Mao, Kitani et al., 2018, Menghini, Corridoni et al., 2019, Moschen, Tilg et al., 2019, Shouval, Biswas et al., 2016). Our previous work as well as that of others has demonstrated that autophagy regulates the production of the proinflammatory cytokines IL-1α, IL-1β and IL-23, primarily in infection models (Castillo, Dekonenko et al., 2012, Dupont, Jiang et al., 2011, Peral de Castro, Jones et al., 2012, Reed, Morris et al., 2015, Watson, Manzanillo et al., 2012, Zhang, Kenny et al., 2015). This observation is the canonical pathway frequently described linking autophagy dysregulation to excess inflammation via IL-1 and IL-17. IL-1 and IL-23 are key regulators of IL-17-mediated inflammation (Chung, Chang et al., 2009, McGeachy, Chen et al., 2009). However, the role of IL-17 in IBD pathogenesis has recently been questioned as therapeutic targeting of IL-17 exacerbates inflammation in both IBD patients and animal models of IBD (Fujino, Andoh et al., 2003, Hueber, Sands et al., 2012, Ogawa, Andoh et al., 2004, Targan, Feagan et al., 2016, Yang, Chang et al., 2008), suggesting that IL-17 may not be the predominant driver of autophagy-linked IBD pathogenesis. Thus, there is a clear gap in knowledge regarding which inflammatory pathway might underlie IBD pathology with respect to autophagy dysregulation.

This study assessed the role of the autophagy gene *Atg5* in myeloid cells in maintaining the balance between intestinal homeostasis and inflammation. Atg5’s most well-understood actions are via the Atg5-Atg12-Atg16L1 complex, which acts as the E3 enzyme conjugating PE to LC3, and along with the E1-like actions of Atg7 and the E2-like actions of Atg3, this pathway drives isolation membrane formation and eventual autophagosome maturation (Hanada, Noda et al., 2007, Mizushima, Noda et al., 1998, Noda, Fujioka et al., 2013, Sakoh-Nakatogawa, Matoba et al., 2013). Atg5 is also embedded in autophagosomal membranes which allows it to interact with fusion proteins in lysosomal membranes such as Tectonic β-propeller repeat containing 1 (TECRP) that facilitate autophagosome-lysosome fusion (Chen, Fan et al., 2012, Ye, Zhou et al., 2018). Thus, Atg5 plays a major role in selective and bulk autophagy which are critical for cell-autonomous immunity. However, it is unclear how Atg5 expression, specifically in myeloid cells, functions outside of bacterial recognition and processing. Here, we show at steady state that mice with an *Atg5*-deficiency in myeloid cells (herein called Atg5ΔMye mice) (Castillo et al., 2012, Zhao, Fux et al., 2008) show alterations in the gut microbiota as well as mucosal TH1 (IFNγ) skewing. Atg5ΔMye mice were susceptible to chemically induced colitis that was characterized by an enhanced IFNγ response. Both TH1 skewing and microbiota changes were partly driven through IL-12 dysregulation in *Atg5*-deficient myeloid cells. Confirming our findings, Atg5ΔMye mice crossed to *IL12p35*-deficient mice resulted in restoration of the gut microbiota and protection from dextran sodium sulfate (DSS)-induced colitis through the reduction of the IL-12 pathway. We further show IL-12 regulation was not mediated through autophagy but interactions with sequestosome-1-like receptor NBR1 and Atg5 action on late endosomes (LE). There has not been a single description of autophagic proteins regulating IL-12-driven immune responses. Thus, these data indicate a new autophagy-independent role for Atg5 and NBR1 in myeloid cells in influencing intestinal homeostasis through an IL-12 pathway.

## RESULTS

### Altered colonic microbiota in mice with an *Atg5*-deficiency in myeloid cells

The gut microbiota is influenced by environmental factors, immune responses, and genetics which is highlighted in individuals with IBD. Numerous studies have reported a decrease in bacterial diversity in IBD patients with major alterations in the phyla Firmicutes and Bacteroidetes (Imhann, Vich Vila et al., 2018, Kang, Alvarado et al., 2020, Manichanh, Borruel et al., 2012, Manichanh, Rigottier-Gois et al., 2006, Ni, Wu et al., 2017, Willing, Dicksved et al., 2010). There is substantial evidence that intestinal myeloid cells are regulated by commensal microbiota (Bain, Bravo-Blas et al., 2014, Chang, Hao et al., 2014, Kang et al., 2020, Mortha, Chudnovskiy et al., 2014, Singh, Gurav et al., 2014); however, numerous studies have emerged demonstrating myeloid regulation of the microbiota (Bader, Enos et al., 2018, Niess, Brand et al., 2005, Rescigno, Urbano et al., 2001, Vallon-Eberhard, Landsman et al., 2006). Autophagy genes appear to be involved in the latter as it was shown the autophagy protein, Atg16L1, expressed in myeloid cells from both humans and mice altered IgA-coated intestinal bacteria at steady-state and during inflammation (Zhang et al., 2017); however, it was not confirmed if this increase in IgA-coated bacteria affects the overall microbial community. More recently, another mouse model with a global knockin of the IBD risk allele *ATG16L1 T300A* had an altered gut microbiota prior to colitis induction (Lavoie et al., 2019).

Utilizing Atg5ΔMye mice (myeloid specific loss of autophagy related 5, *Atg5*, gene), we investigated the importance of Atg5 expression in myeloid cells in maintaining the intestinal microbiota (Castillo et al., 2012, Zhao et al., 2008). In comparison to littermate controls (Atg5-Wild-type, Atg5-Wt), Atg5ΔMye mice showed significant differences in the intestinal microbial composition (Fig. 1 A). Similar to what is observed in IBD patients (Manichanh et al., 2012, Manichanh et al., 2006, Ni et al., 2017, Seksik, 2010, Willing et al., 2010), Atg5ΔMye mice had a decrease in the phylum Firmicutes (Fig. 1 B) and an increase in the phylum Bacteroidetes (Fig. 1 C) compared to Atg5-Wt mice. No significant changes were observed for phyla Actinobacteria and Proteobacteria (Fig. 1, D and E). Additionally, the principal coordinate analysis (PCoA) plot and heatmap of bacterial communities revealed tight clustering of Atg5-Wt microbiota that was distinct from Atg5ΔMye microbiota at the genus level (Fig. 1, F and G) and this was also observed for female mice (Supp. Fig. 1 A and B). Thus, Atg5 expression in myeloid cells is critical for the maintenance of the intestinal microbiota.

**Figure 1.**
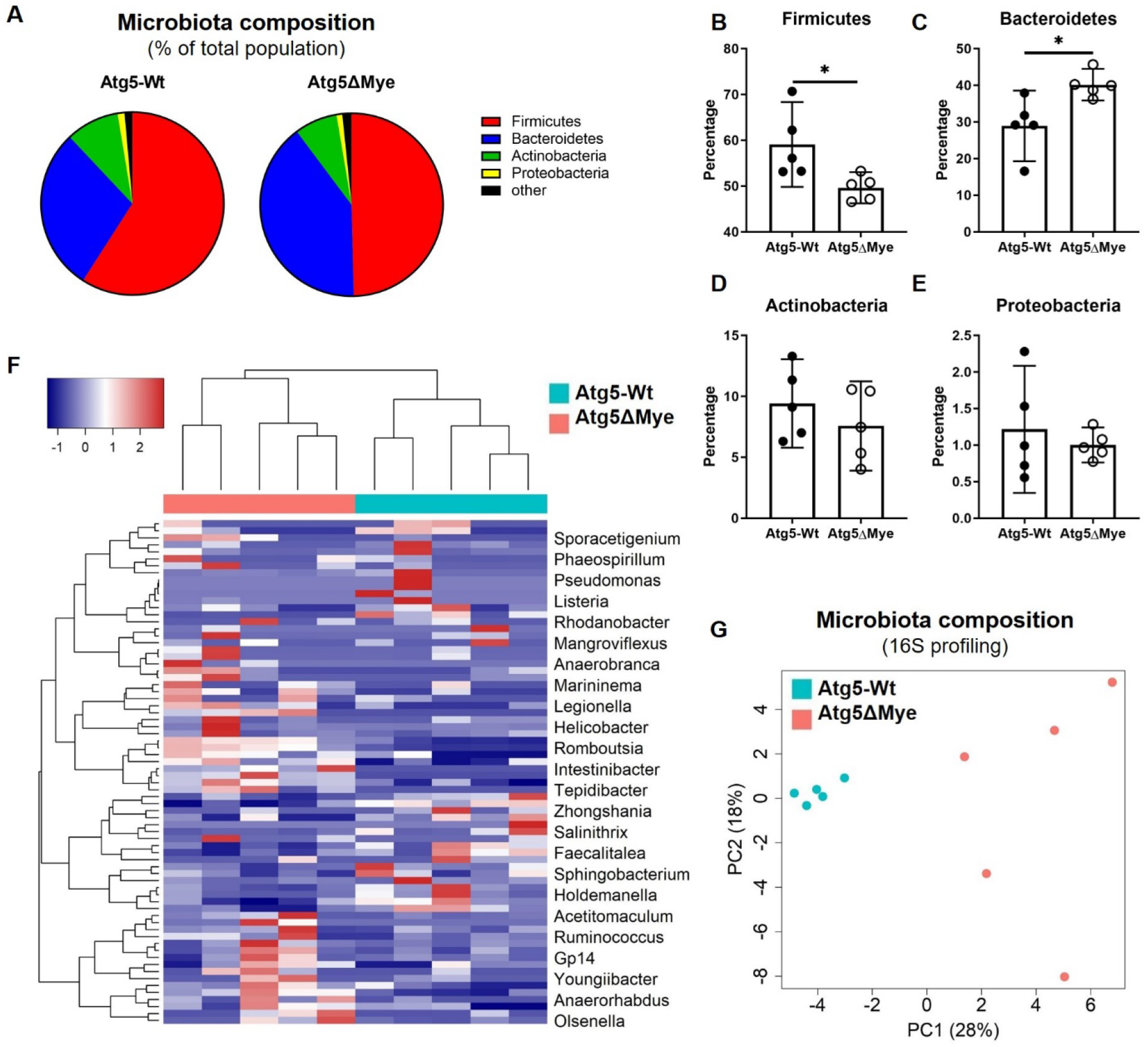
Alterations in the composition of the colonic microbial community in mice with a selective *Atg5*-deficiency in myeloid cells. Bacterial composition as assessed by 16S rRNA sequencing of DNA extracted from freshly collected stool of Atg5-Wt and Atg5ΔMye mice (n=5 per group). **(A)** Pie chart showing the average proportion of Firmicutes, Bacteroidetes, Actinobacteria, Proteobacteria and all other phyla in 8-week old mice. **(B-E)** Quantification of the phyla Firmicutes, Bacteroidetes, Actinobacteria, and Proteobacteria. **(F)** Heatmap of the relative abundance of colonic microbes (Genus level). **(G)** Principal coordinates analysis (PCoA) plot of microbiota composition (Genus level). Data is shown as mean (±95%CI) and a T-test was used to measure specific microbiome species abundance between groups. Adjusted p-value > 0.05 was used as significant threshold.

### Myeloid Atg5 expression regulates IFNγ response in the intestinal microenvironment

The proinflammatory cytokine IL-17 contributes to shaping and regulating the intestinal microbiota. We and others have shown T cells from mice wherein myeloid cells lack *Atg5* or other autophagy-related genes display skewing towards IL-17 polarization (Castillo et al., 2012, Peral de Castro et al., 2012, Reed et al., 2015, Watson et al., 2012). This T cell polarization is mediated through the dysregulation of cytokines or cellular components that promote IL-17 responses (Chung et al., 2009, McGeachy et al., 2009). Microbiota differences can also alter T cell polarization (Atarashi, Tanoue et al., 2011, Chen, Sun et al., 2019, Ivanov, Atarashi et al., 2009, Ivanov, Frutos Rde et al., 2008, Sun, Wu et al., 2018). To determine whether an enhanced level of IL-17-producing T cells populating the intestinal mucosa is influencing the microbiota of Atg5ΔMye mice, we isolated and stimulated CD4^+^ T cells from the mesenteric lymph nodes (mLN) to determine TH polarization. Interestingly, we observed an increase in IFNγ-producing CD4^+^ T cells from Atg5ΔMye mice (Supp. Fig. 1 C,D) with no significant difference in the number of IL-17-producing CD4^+^ T cells (Supp. Fig. 1 E). Thus, at steady state, Atg5ΔMye mice show differences in CD4^+^ T cell polarization.

To examine the impact of TH1 skewing and microbiota alteration in Atg5ΔMye mice during an inflammatory state, we subjected both Atg5ΔMye and Atg5-Wt mice to DSS-induced acute colitis. Similar to what was reported in mice with an *Atg16/1*- or *Atg7*-deficiency specifically in myeloid cells (Lee, Kim et al., 2016a, Zhang et al., 2017), colitis induction in Atg5ΔMye mice resulted in an exacerbated inflammatory response compared to Atg5-Wt mice. Intestinal epithelial disruption via DSS caused significant weight loss (Fig. 2 A) as well as acute diarrheal response and mucosal bleeding (disease activity index, DAI, Fig. 2 B) in Atg5ΔMye mice. The length of the colon was measured to assess colonic mucosal injury and revealed colonic shortening and severe inflammation from the cecum to the distal colon of Atg5ΔMye mice (Fig. 2 C and Supp. Fig. 1, F and G). Additionally, we found that the spleen weight of colitic Atg5ΔMye mice was increased (Fig. 2 D). These results indicate Atg5 expression in myeloid cells, like Atg7 and Atg16L1, is critical to control acute intestinal inflammation.

**Figure 2.**
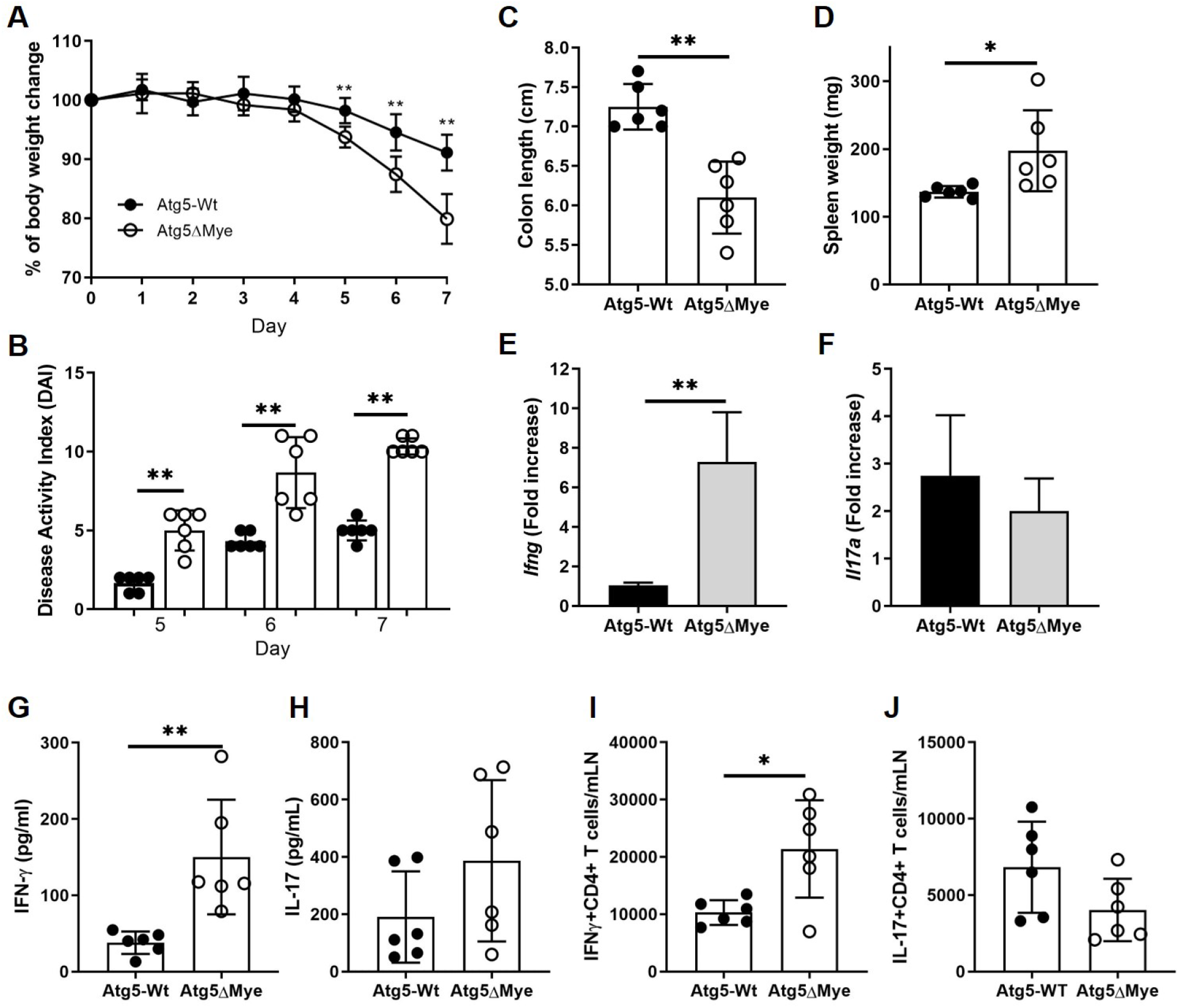
Myeloid Atg5 expression prevents excessive IFNγ-mediated intestinal inflammation. DSS-induced colitis and T cell response in Atg5-Wt and Atg5ΔMye mice. **(A)** Percent weight loss between Atg5-Wt and Atg5ΔMye mice. **(B)** Disease activity index (DAI) as determined by weight loss, behavior, acute diarrheal response and mucosal bleeding. **(C)** Colon length after colitis. **(D)** Spleen weight after colitis. **(E and F)** Colonic *Ifng* and *Il17a* gene expression after colitis induction. **(G and H)** Colonic IFNγ and IL-17 protein expression as determined by ELISA. **(I and J)** Percent of IFNγ^+^ and IL-17^+^ CD4^+^ T cells isolated from the mLN of colitic mice. Representative of two independent experiments, Graphs indicate mean (±SD). * P<0.05, **P<0.01. Two-tailed unpaired Student’s t tests or by 2-way ANOVA with Tukey’s post hoc test.

The acute DSS model is an innate/wound repair model and although the acute DSS model is not solely dependent on B and T cell responses, both participate in the exaggerated presentation of the disease. In fact, T cells have been shown to be the major driver of colonic inflammation in the acute DSS model by day 4 with a peak at day 8 (Nunes, Kim et al., 2018). Interestingly, we do not see major changes in DAI in Atg5ΔMye mice until day 6 and 7 (Fig. 2B) when we would expect T cells to be responding. Furthermore, IFNγ has been shown to play an indispensable role in the initiation of acute DSS colitis and IFNγ-deficient mice are protected from acute DSS induced colitis (Ito, Shin-Ya et al., 2006). Thus, this model is sufficient to show IFNγ responses from T cells in Atg5ΔMye mice. Our observation that CD4^+^ T cells are skewed towards a type 1 immune response in Atg5ΔMye mice at steady state prompted an examination of cytokine expression and TH skewing during colitis. Colitic colons from Atg5ΔMye mice showed increased IFNγ gene and protein expression compared to controls (Fig. 2, E and G). Similar to steady state, there was also an increase in IFNγ-producing CD4^+^ T cells isolated from the mLN of colitic Atg5ΔMye mice (Fig. 2 I). While IL-17 was expressed in both Atg5ΔMye and Atg5-Wt mice, we found no significant difference in IL-17 expression in the colon (Fig. 2, F and H) or from mLN CD4^+^ T cells (Fig. 2 J). This discrepancy in IL-17 expression could be tissuespecific as models in which myeloid cells lack autophagic components and express high levels of IL-17 have been reported for respiratory infections (Castillo et al., 2012, Reed et al., 2015, Watson et al., 2012). However, others have reported TH1 skewing in ATG16L1T300A knockin mice as well as in conditional myeloid Atg7-deficient mice during intestinal inflammation (Lavoie et al., 2019, Zhang et al., 2017). Thus, our data suggest Atg5 expression in myeloid cells protects the intestinal microenvironment during inflammation through limiting IFNγ expression.

### Atg5 limits IL-12 production in myeloid cells independent of canonical autophagy

The current paradigm is that an autophagic defect (or loss of an autophagy-related gene) in myeloid cells leads to enhanced IL-1α/β expression (Castillo et al., 2012, Dupont et al., 2011, Peral de Castro et al., 2012). The excess production of IL-1 skews lymphocytes to produce IL-17, subsequently promoting IL-17-mediated inflammation. However, we found colitic Atg5ΔMye mice presented with increased IFNγ and no significant difference in IL-17 in the colon microenvironment. This suggests colonic *Atg5*-deficient myeloid cells are likely to produce other factors that promote TH1 skewing and IFNγ production. IL-12p70 (hereafter called IL-12) is a major cytokine involved in TH1 polarization and is a potent inducer of IFNγ (Eftychi, Schwarzer et al., 2019, Moschen et al., 2019, Perussia, Chan et al., 1992, Trinchieri, Wysocka et al., 1992). IL-12 is a heterodimeric protein comprised of IL-12p35 and IL-12p40 (IL-12p40 also dimerizes with IL-23p19 to form IL-23, a major cytokine maintaining TH17 cells) and is highly expressed by various myeloid cells including macrophages (Abdi & Singh, 2015, Carra, Gerosa et al., 2000, D’Andrea, Rengaraju et al., 1992, Moschen et al., 2019, Reitberger, Haimerl et al., 2017). We have previously shown Atg5ΔMye mice also produce excess IL-12 during tuberculosis infection (Castillo et al., 2012). Utilizing bone marrow-derived macrophages (BMM) from Atg5ΔMye and Atg5-Wt mice, we showed that *Atg5*-deficient macrophages secrete excess IL-12 upon LPS/IFN-γ stimulation compared to Atg5-Wt macrophages (Fig. 3A). For many cell types, low levels of IL-12p35 are constitutively expressed, while IL-12p40 expression occurs primarily in macrophages and dendritic cells, and both increase in response to microbial stimulation (Abdi & Singh, 2015, Carra et al., 2000, D’Andrea et al., 1992, Reitberger et al., 2017). We observed no difference in *Il12a* (IL-12p35) and *Il12b* (IL-12p40) gene expression (Supp. Fig. 2, A and B). Furthermore, there was no difference in cell surface expression of IFNγ-receptor, TLR4/MD2 or CD14 between Atg5-Wt and *Atg5*-deficient BMM (Supp. Fig. 2 C) that would enable an enhanced response. This data suggests Atg5 has a post-transcriptional role in regulating IL-12 secretion.

**Figure 3.**
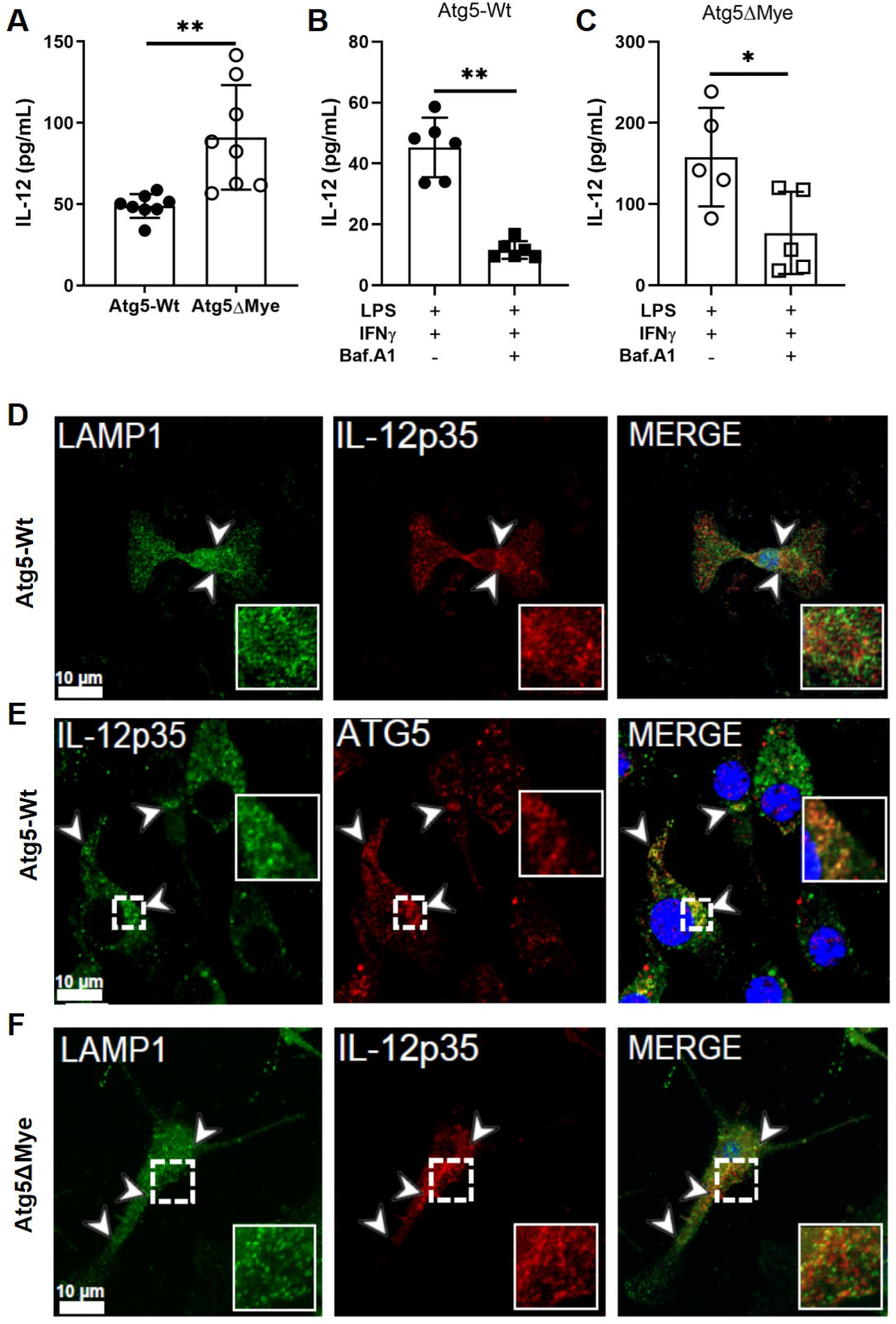
Atg5 regulates IL-12 secretion in macrophages. Analysis of IL-12 secretion in macrophages. **(A)** Detection of IL-12 by ELISA from Atg5-Wt or Atg5ΔMye BMM after LPS and IFNγ stimulation. **(B and C)** Detection of IL-12 by ELISA from Atg5-Wt **(B)** or Atg5ΔMye **(C)** BMM after LPS and IFNγ stimulation in the presence or absence of Baf. A1. Representative of two independent of experiments. **(D - F)** 1-3 μm Z-stack images were performed on BMM stimulated with LPS, IFNγ and Baf. A1 using immunofluorescence at 1.6 Zoom by a 63X oil immersion objective. **(D)** Atg5-Wt BMM were stained for LAMP1 (green) and IL-12p35 (red). **(E)** Atg5-Wt BMM were stained for ATG5 (red) and IL-12p35 (green). **(F)** Atg5ΔMye BMM were stained for LAMP1 (green) and IL-12p35 (red). Representative from 65 images from 15 slides from two independent of experiments. Arrows in images indicate puncta co-localizing with insets displaying enlargement of indicated region. Scale bars: 10μm. Graphs indicate mean (±SD). * P<0.05, **P<0.01,***P<0.001. Two-tailed unpaired Student’s t tests.

As mentioned above, Atg5 assists in autophagosome formation and autophagosome-lysosome fusion. Thus, the genetic deletion of *Atg5* would consequently affect several steps in autophagy. We next considered whether IL-12 was a direct target for autophagic removal. However, endogenous IL-12 and LC3 did not colocalize during stimulation and treatment with bafilomycin A1 (Baf. A1) (Supp. Fig. 2 D) in Atg5-Wt or *Atg5*-deficient BMM suggesting IL-12 is unlikely to be a direct target for autophagic degradation. Baf. A1 is widely used to inhibit autophagic flux by targeting the V-ATPase ATP6V0C/V0 subunit c but can also deacidify endosome/lysosome vesicles through the same mechanism (Harada, Shakado et al., 1997, Yoshimori, Yamamoto et al., 1991). Atg5 also regulates acidification and deacidification of late endosomal compartments (Guo, Chitiprolu et al., 2017) and prior to stimulation *Atg5*-deficient BMM display increased lysotracker staining compared to Atg5-Wt BMM (Supp Fig. 2E) suggesting the loss of Atg5 affects the regulation of vesicles’ pH levels (Guo et al., 2017, Peng, Zhang et al., 2014). Interestingly, Baf. A1 treatment decreased IL-12 secretion in Atg5-Wt BMM (Fig. 3 B). Furthermore, treatment of *Atg5*-deficient BMM with Baf. A1 also reduced IL-12 secretion (Fig. 3 C). These data suggest Atg5 regulates IL-12 secretion through vesicle acidification and Baf. A1 can compensate for the loss of Atg5 in regulating IL-12 secretion.

We next examined the intracellular localization of IL-12 after Baf. A1 treatment. As mentioned above, IL-12 consists of subunits IL-12p35 and IL-12p40. While the IL-12p40 subunit has conventional secretory sequences and can be secreted in its homodimeric form, the IL-12p35 peptide is leaderless, cannot be secreted as a monomer, and conventional secretion cannot be induced by the addition of a secretory sequence (Abdi & Singh, 2015, Carra et al., 2000, Reitberger et al., 2017). The purpose of IL-12p35’s constitutive expression remains poorly understood (Abdi & Singh, 2015) but a recent report demonstrated IL-12p35 is trafficked to late endosomes (LE) before secretion (Chiaruttini, Piperno et al., 2016). As observed with total IL-12, Baf. A1 treatment resulted in IL-12p35 accumulation in LE as determined by IL-12p35 (red) and Lamp1 (green) colocalization (Fig. 3 D). White arrows in images indicate puncta colocalizing with insets displaying enlargement of indicated region. Interestingly, IL-12p35 (green) colocalized with Atg5 (red) (Fig. 3 E). IL-12 (red) colocalization with Lamp1 (green) was more evident in *Atg5*-deficient BMM (Fig. 3 F). Further analysis, revealed IL-12 (red) also colocalized with the LE marker Rab7 (green) (Supp. Fig. 2F) as previously reported (Chiaruttini et al., 2016). Taken together, our data suggest Lamp1^+^ LEs (Cheng, Xie et al., 2018, Dunster, Toh et al., 2002) contain IL-12 and the lack of Atg5 leads to increased levels of IL-12 in these vesicles.

### The sequestosome-1-like receptor NBR1 is involved in IL-12 secretion

While most cytokines are directed by their signal sequence through endoplasmic reticulum (ER)-golgi complex pathway for processing and trafficking, some inflammatory cytokines, including IL-1β and IL-18, are known to be excreted via alternative strategies (Abdel Fattah, Bhattacharya et al., 2015, Claude-Taupin, Bissa et al., 2018, Dupont et al., 2011, Kimura, Jia et al., 2017, Murai, Okazaki et al., 2015). For IL-1β, the selective autophagy cargo receptor TRIM16 directs IL-1β to LC3-II+ sequestration membranes for secretion (Kimura et al., 2017).

This suggests other selective cargo receptors such as the sequestosome-1-like receptors (SLRs: p62/SQSTM1 and NBR1) could act as possible cargo receptors for alternative secretion (Claude-Taupin, Jia et al., 2017, Jiang, Dupont et al., 2013). We considered NBR1 might be responsible for delivering IL-12 to LE for secretion given a homolog of the mammalian NBR1, Nbr1 from *Schizosaccharomyces pombe* was shown to deliver proteins to LE (Mardakheh, Yekezare et al., 2009) and NBR1 accumulates upon autophagy inhibition (Kirkin, Lamark et al., 2009). Indeed, confocal imaging revealed NBR1 (red) co-localized with IL-12p35 (green) in WT BMM after LPS/IFN-γ and Baf. A1 treatment (Fig. 4 A, top row and B); white arrows indicate puncta co-localizing. However, this co-localization was reduced upon brefeldin A (BrefA) treatment, suggesting NBR1 (red) and IL-12 (green) interaction occurs after IL-12 leaves the ER (Fig. 4 A, bottom row and B). This reduction in colocalization was not due to enhanced secretion as IL-12 was undetectable by ELISA after LPS/IFNγ/brefA treatment (data not shown). To verify NBR1/IL-12 interaction in macrophages, cell lysates were immunoprecipitated with anti-IL12p35, anti-NBR1 or isotype control antibodies and subjected to western blot with anti-IL-12p35, anti-IL-12p40, anti-NBR1 and anti-p62 antibodies. Both IL-12p40 and NBR1 coprecipitated with anti-IL-12p35 antibodies (Fig. 4 C). Reciprocally, IL-12p35 and IL-12p40 coprecipitated with anti-NBR1 antibodies (Fig. 4 C); however, neither probe co-precipitated p62/SQSTM1 (Fig. 4C). This interaction was further verified *in silico* through ClusPro docking (Kozakov, Hall et al., 2017) as IL-12 (1F45) interacted with multiple peptide fragments (1WJ6, 2L8J, 2MGW, and 2MJ5) of NBR1 (Fig. 4 D and E). Lastly, a knock down of NBR1 via small interfering RNA (siRNA) in Atg5-Wt BMM reduced IL-12 secretion compared to scrambled siRNA control (Fig. 4 F, Supp. Fig. 2 G). A reduction in IL-12 secretion was also observed in *Atg5*-deficient BMM after NBR1 knockdown suggesting this interaction is independent of Atg5 (Fig. 4 G, Supp. Fig. 2 G). Taken together, our data suggest NBR1 is partially responsible for IL-12 secretion whereby it functions to direct IL-12p35 to LE for secretion.

**Figure 4.**
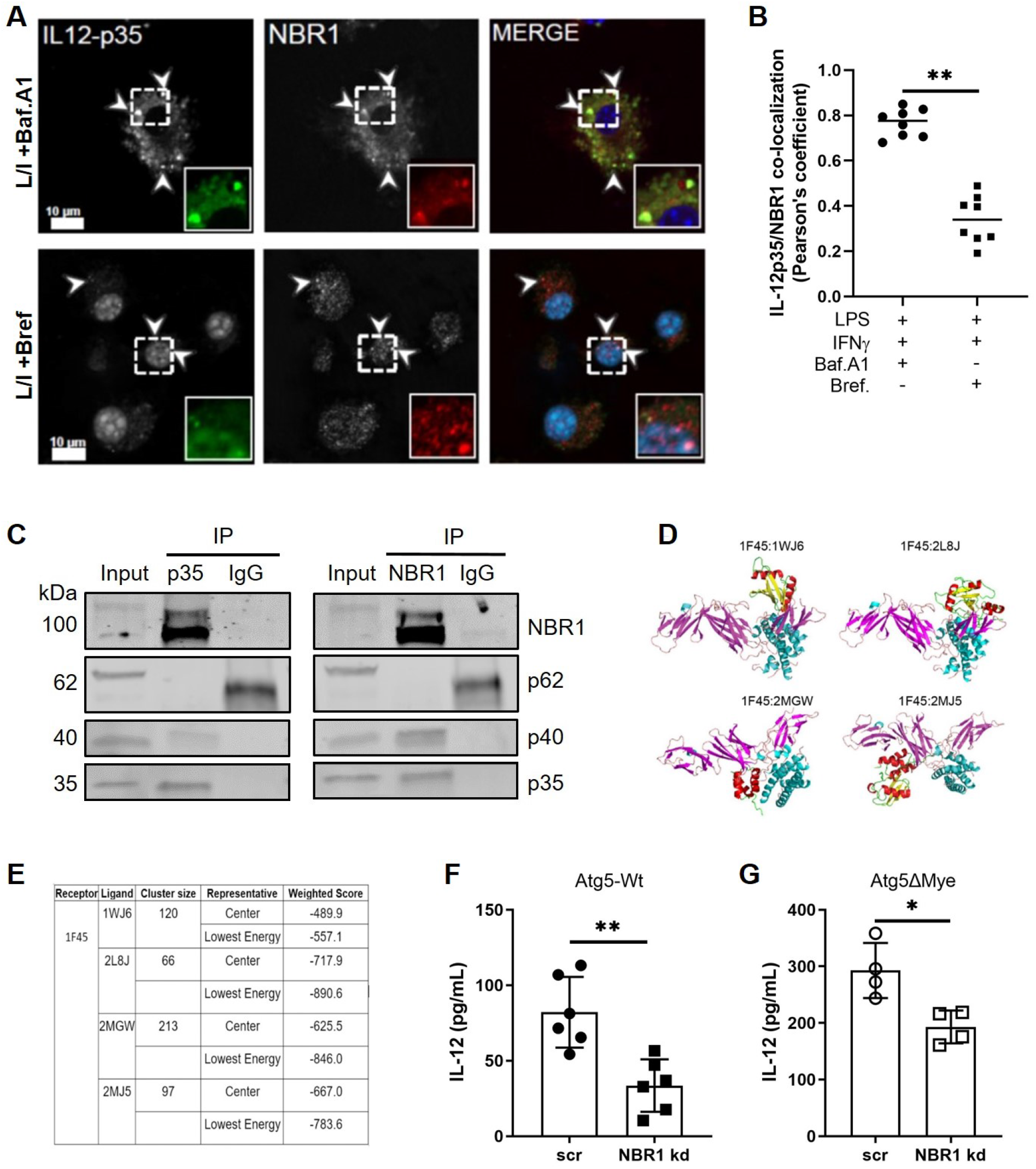
NBR1 and IL-12 interaction. Analysis of NBR1 and IL-12 interaction in macrophages. **(A)** Atg5-Wt BMM were stained for NBR1 (red) and IL-12p35 (green) after LPS and IFNγ stimulation and the addition of Bafilomycin A1 (Top Row) or Brefeldin A (Bottom Row). **(B)** Quantification was performed using Pearson’s correlation coefficient (co-localization) using image analysis. Representative of two independent of experiments and of 10 images from 5 slides. Arrows in images indicate puncta co-localizing with insets displaying enlargement of indicated region. Scale bars: 10μm. **(C)** Anti-IL-12p35 antibody co-immunoprecipitated NBR1 and IL-12p40, and anti-NBR1 antibody co-immunoprecipitated IL-12p35 and IL-12p40. **(D and E)** Protein docking between IL12A (PDB ID: 1F45) and NBR1. **(D)** the docking models and **(E)** ClusPro docking results for four candidate interaction models between 1F45 and four peptide fragments of NBR1 (PDB IDs: 1WJ6, 2L8J, 2MGW, and 2MJ5). **(F and G)** Effects of NBR1 knockdown on IL-12 secretion in Atg5-Wt or Atg5ΔMye BMM. Graphs indicate mean (±SD). * P<0.05, **P<0.01. Two-tailed unpaired Student’s t tests.

### Limiting IL-12 secretion restores the gut microbiota and protects against DSS-induced colitis in mice with an *Atg5*-deficiency in myeloid cells

To determine if the change in the gut microbiota and susceptibility to DSS-induced colitis in Atg5ΔMye mice was due to the excess IL-12 secretion, we crossed Atg5ΔMye mice to IL-12p35 deficient mice (Mattner, Magram et al., 1996) to generate Atg5ΔMye-IL12KO mice. We first examined the colonic microbiota at steady state in all three groups of mice (i.e. Atg5-Wt, Atg5ΔMye and Atg5ΔMye-IL12KO). Atg5ΔMye-IL12KO mice showed a global change in the intestinal microbial composition in comparison to Atg5ΔMye mice, particularly an increase in the phyla Firmicutes (Supp. Fig. 3 A). Furthermore, the PCoA plots and heatmap show clustering of the bacterial communities (genus level) of the Atg5ΔMye-IL12KO and Atg5-Wt colonic mice that was separated from the bacterial community in the Atg5ΔMye mice (Fig. 5, A and B).

**Figure 5.**
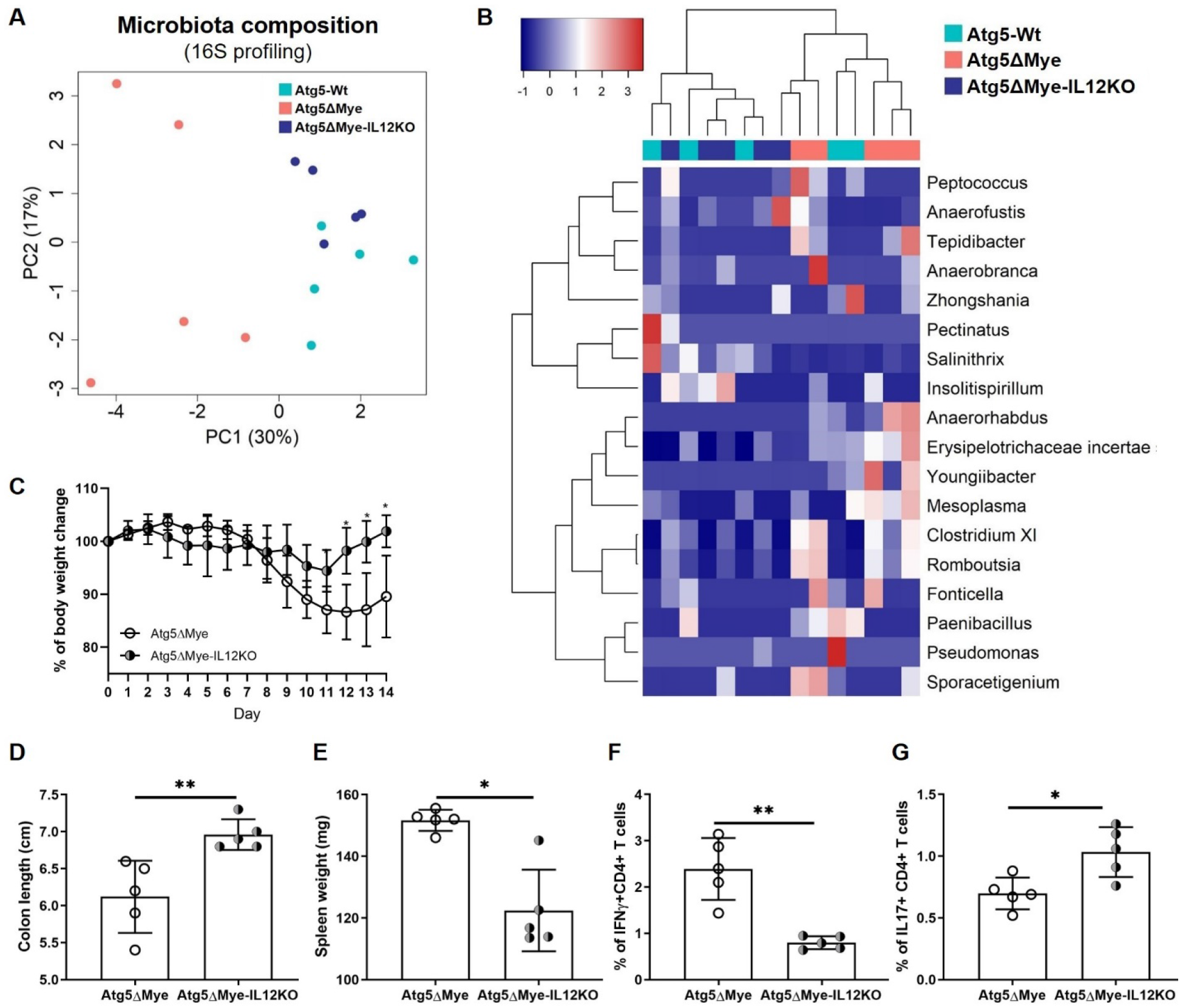
Limiting IL-12 in mice with a selective *Atg5*-deficiency in myeloid cells restores the intestinal microbiota and alleviates inflammation. Bacterial composition as assessed by 16S rRNA sequencing from freshly collected stool of Atg5ΔMye and Atg5ΔMye-IL12KO mice prior to colitis induction (n=5 per group). **(A)** Principal coordinates analysis (PCoA) plot of microbiota composition (Genus level). **(B)** Heatmap of the relative abundance of colonic microbes (Genus level). A T-test was used to measure specific microbiome species abundance between groups. Adjusted p-value > 0.05 was used as significant threshold.. **(C)** Percent weight loss between Atg5ΔMye and Atg5ΔMye-IL12KO mice. **(D)** Colon length after colitis. **(E)** Spleen weight after colitis. **(F and G)** Percent of IFNγ^+^ and IL-17^+^ CD4^+^ T cells isolated from the mLN of colitic mice. Graphs indicate mean (±SD). * P<0.05, **P<0.01. Two-tailed unpaired Student’s t tests or by 2-way ANOVA with Tukey’s post hoc test.

We next examined Atg5ΔMye-L12KO mice response to DSS-induced colitis. Atg5ΔMye-IL12KO mice were protected from DSS-induced colitis compared to Atg5ΔMye mice. Atg5ΔMye-IL12KO mice suffered minimal weight loss, showed no shortening of the colon after colitis induction, and had reduced spleen weights (Fig. 5, C-E). There was also a decrease in *Ifng* gene expression in the colon after colitis induction in Atg5ΔMye-IL12KO mice but no change in *Il17a* gene expression between both groups of mice. (Supp. Fig. 3, B and C). Additionally, we observed a decrease in the percent of IFNγ-producing CD4^+^ T cells from the mLN of Atg5ΔMye-IL12KO mice compared to Atg5ΔMye mice (Fig. 5 F). However, Atg5ΔMye-IL12KO mLN CD4^+^ T cells did produce more IL-17 (Fig. 5 G). This change in IL-17 expression is likely due to *Atg5*-deficient myeloid cells producing excess IL-1 (Castillo et al., 2012). Nevertheless, genetic deletion of IL-12p35 from mice in which myeloid cells also lack Atg5 restores the microbiota and protects against DSS-induced colitis.

## DISCUSSION

This work identifies a role for Atg5 in myeloid cells in unconventional cytokine secretion that consequently affects intestinal homeostasis. Additionally, it adds to the plethora of functions described for Atg5 (Guo et al., 2017, Inomata, Into et al., 2013, Lee, Mattei et al., 2010, Ndoye, Budina-Kolomets et al., 2017, Peng et al., 2014, Simon, Yousefi et al., 2014, Yousefi, Perozzo et al., 2006). Along with these studies, this work establishes a new non-autophagy, immunological role for Atg5 in promoting TH1 responses. In the context of IBD and autophagy-related proteins (Atg), GWAS has only identified *ATG16L1* variants (Hampe et al., 2007). Nevertheless, animal models with an Atg5-deficiency display a similar phenotype to Atg16L1-deficient mice (Cadwell et al., 2009). Interestingly, the levels of *ATG5* and the function of autophagy (and possibly other ATG5 functions) are decreased in IBD patient samples. This decrease in *ATG5* expression and autophagy activity in IBD patients is linked to an increased expression of the microRNA miR30C that acts to downregulate *ATG5* expression (Nguyen, Dalmasso et al., 2014, Ye et al., 2018). Therefore, inhibiting Atg5 expression and function, described here in only myeloid cells, ultimately alters the intestinal microenvironment.

As previously reported, *Atg5*-deficient myeloid cells produce excess proinflammatory cytokines including IL-1α and IL-1β (Castillo et al., 2012, Dupont et al., 2011). Thus, we cannot dismiss that either cytokine contributes to the excess IFNγ observed in Atg5cKO mice as IL-1 can synergize with IL-12 to enhance IFNγ production and TH1 skewing (Cooper, Fehniger et al., 2001, Tominaga, Yoshimoto et al., 2000). Although we and others did not find any changes in colonic IL-17-producing cells in mice with a genetic deletion of Atg5 or Atg7 in myeloid cells (Castillo et al., 2012, Lee et al., 2016a), both colonic TH17 and TH1 cells were found to be increased in *ATG16L1* T300A knockin mice (Lavoie et al., 2019). Given Atg5ΔMye-IL12KO mice were protected against colitis induction, these results together suggest IL-12 is a key cytokine driving intestinal pathology and TH1 skewing in our model.

This similarilty in intestinal TH1 responses between these mouse models is likely linked to the functional role of Atg5, Atg7 and Atg16L1. All three proteins are involved in some aspect of the autophagy process as well as endosome and lysosome regulation (Fraser, Simpson et al., 2019, Galluzzi et al., 2017, Mardakheh et al., 2009). The role of the TH17 response in driving intestinal pathology is up for debate as there is strong evidence suggesting IL-17 may have a protective role in the intestine. Exacerbated intestinal inflammation was reported in *Il17a*-deficient mice or after *in vivo* blockade of IL-17 during DSS-induced colitis (Ogawa et al., 2004, Yang et al., 2008). Furthermore, in clinical trials for IBD patients, targeting IL-17 or IL-17R worsened symptoms leading to early clinical trial termination (Hueber et al., 2012, Targan et al., 2016). Nevertheless, this does not rule out the effects of increased levels of IL-17 during intestinal inflammation (Fujino et al., 2003, Jiang et al., 2014, Moschen et al., 2019). In fact, IL-17 plays a major role in regulating mucosal IgA production (Dann, Manthey et al., 2015, Kumar, Monin et al., 2016). Interestingly, increased IgA-coated bacteria were found in mice that lack Atg16L1 in myeloid cells as well as in the stool of Crohn’s disease patients who were homozygous for the *ATG16L1* T300A risk allele (Zhang et al., 2017). *ATG16L1* T300A knockin mice also showed alteration in the microbiota. Similar to our findings, *ATG16L1* T300A knockin mice also had a decrease in the phyla Firmicutes and an increase in the phyla Bacteroidetes (Lavoie et al., 2019). It is unclear if these microbiota changes in *ATG16L1* T300A knockin mice are due specifically to myeloid cells, since all cells including the intestinal epithelium express the *ATG16L1* T300A gene in this mouse model. Additionally, deletions or polymorphisms of autophagy genes in myeloid cells alter bacterial clearance suggesting there are multiple mechanisms by which autophagy genes regulate host-microbiota interactions. Our data suggest myeloid cells likely have a strong contribution in maintaining the microbiota through Atg5’s action on IL-12 secretion.

The autophagic process and the genes that regulate autophagy are crucial for intestinal homeostasis. Autophagy is required for the maintenance of tight junction integrity, gut-commensal homeostasis, and control of invasive bacteria at the intestinal epithelium (Bel et al., 2017, Benjamin et al., 2013, Burger et al., 2018, Cadwell et al., 2008, Cadwell et al., 2009, Lee et al., 2016a, Matsuzawa-Ishimoto et al., 2017, Nighot et al., 2015, Patel et al., 2013, Wong et al., 2019, Wu, Wang et al., 2017). Multiple lines of evidence also suggest autophagy regulates inflammatory cytokines (Castillo et al., 2012, Dupont et al., 2011, Lee, Foote et al., 2016b, Merkley, Chock et al., 2018, Saitoh, Fujita et al., 2008). These attributes make modulation of autophagy or single autophagic proteins an excellent therapeutic target. Nevertheless, it is critical to understand the function of these individual proteins given they can modulate different TH responses. In fact, reagents that are known to target autophagy activation such as rapamycin, increase IL-12 secretion (Macedo, Turnquist et al., 2013, Ohtani, Nagai et al., 2008). Additionally, chloroquine, a known inhibitor of autophagosome-lysosome fusion, reduces IL-12 secretion (Said, Bock et al., 2014), likely through a similar mechanism as Baf. A1. These reagents have been used with some clinical efficacy in IBD patients and animal models of IBD (Goenka, Kochhar et al., 1996, Kanvinde, Chhonker et al., 2018, Nagar, Ranade et al., 2014, Park, Jang et al., 2019). It is unclear if these effects were due to IL-12 regulation but targeting IL-12, specifically, the IL-12p40 subunit that is shared with IL-23, has also proven beneficial for some IBD patients (Feagan, Sandborn et al., 2016, Moschen et al., 2019, Sands, Sandborn et al., 2019). Therefore, understanding how autophagy or single autophagic proteins regulate proinflammatory cytokines will allow precision in modulating the immune system in chronic inflammatory conditions like IBD. Lastly, targeting these autophagy proteins or their functions could be an alternative to mitigate problems associated with biologics including loss of function, immunogenicity and cost (D’Haens, 2007).

In conclusion, our data support a novel role for Atg5 expression in myeloid cells in regulating intestinal homeostasis. Atg5 expression in myeloid cells controls the IL-12-IFNγ pathway that influences the microbiota and limits this pathway during colitis (Supp. Fig. 3 D). Mechanistically, we show both Atg5 and the cargo receptor NBR1 regulate IL-12 secretion in myeloid cells. Specifically, we propose that NBR1 shuttles IL-12 to LE whereby Atg5 functions to control the pH of these IL-12 containing vesicles for secretion. A genetic deletion of Atg5 results in dysregulation of IL-12 secretion as well as the accumulation of SLR’s such as NBR1 (Kirkin et al., 2009) which could potentially allow for increased accumulation of IL-12 in LE. Consequently, these attributes increase IL-12 secretion. A reduction in IL-12 secretion could be achieved by removing NBR1 or reducing LE/lysosomal pH with Baf. A1 even in the absence of Atg5 expression. Nevertheless, the dysregulation of IL-12 in Atg5ΔMye mice leads to alterations of the microbiota and severe colitis when the intestinal barrier is disrupted.

## MATERIALS AND METHODS

### Animals

The transgenic Atg5ΔMye mice (myeloid specific Atg5 deletion) and Atg5-Wt mice have previously been characterized (Zhao et al., 2008). B6.129S1-*Il12a*^tm1jm^/J (IL-12p35-deficient mice) were purchased from JAX (002692) (Mattner et al., 1996) and crossed to Atg5ΔMye mice to generate Atg5ΔMye-IL12KO mice. All mice were genotyped for the presence of *Atg5, Il12a* or Cre expression by Transnetyx Inc. All experiments were approved by the Institutional Animal Care and Use Committee of the University of New Mexico Health Sciences Center, in accordance with the National Institutes of Health guidelines for use of live animals. The University of New Mexico Health Sciences Center is accredited by the American Association ofor Accreditation of Laboratory Animal Care.

### Intestinal inflammation model

For Dextran Sodium Sulfate (DSS)-induced colitis, mice were provided 2.5% DSS (colitis grade, ~30,000 −50,000 MW; MP Biomedicals) in drinking water adlibitum. Reagent used for *in vivo* treatment as well as sample collection and stimulation are included in Table S1.

### Microbiota analysis

Fecal samples from Atg5-Wt, Atg5ΔMye, and Atg5ΔMye-IL-12KO mice were collected fresh in sterile tubes and flash frozen. Microbial communities were determined by sequencing of the 16S rRNA as previously reported (Komesu, Richter et al., 2017) with minor modifications described below. Microbial DNA was isolated from feces using the ZymoBIOMICS DNA Miniprep Kit (Zymo Research) following manufacturer’s recommendations. Variable regions V-3 through V-4 of the 16S rRNA gene were amplified by PCR using 100 ng input of DNA for each sample in duplicate using primers 319F- (5’-ACTCCTRCGGGAGGCAGCAG-3’) and 806R- (5’-GACAGGACTACHVGGGTATCTAATCC-3’) containing Nextera adapter overhangs. A second PCR was performed with Nextera^®^ XT Index Kit v2 Set A (Illumina) to complete the adapter and add a unique sample-specific barcodes. After each PCR, a clean-up with AxyPrep Fragment Select-I magnetic beads (Axygen Biosciences) was completed, and all PCR reactions were run on an Applied Biosystems 2720 Thermal Cycler. Indexed samples were combined to yield duplicate 300 ng pools, followed by the creation and denaturation of a 4 nM library, and paired 250bp sequencing runs were completed on the Illumina MiSeq using v3 sequencing chemistry (Illumina). All reagents and kits are listed in Table S2.

### Microscopy and image analysis

For confocal microscopy, macrophages were plated at 100k cells per well on 18 mm glass coverslips and stimulated with LPS (500 ng/mL) and IFNγ (10 ng/mL) for 8 hours and treated with BrefA (10 nM) or Baf. A1 (10 nM) for 2 hours. Cells were then fixed with 4% PFA followed by a wash with 1x PBS. Blocking buffer contained PBS with 50% FBS, 2% BSA and 0.1% saponin. After a 1-hour stain in primary antibody, cells were washed with PBS then followed by 1-hour stain in blocking buffer containing secondary antibody. Cells were mounted using ProLong Gold Antifade with DAPI and imaged using the Zeiss LSM 800 Airyscan Confocal microscope with a 63x oil objective lens. Images were processed using Zen Software and Adobe Photoshop (version CC 2019). All primary and secondary antibodies used for confocal staining are listed in Table S3.

### Statistical Analysis

Statistical analysis was performed as described in figure legends and graphs generated display mean (± SD) and were obtained using Prism software. Microbiome data was sequenced and processed by Illumina’s service lab using their inhouse analysis pipeline. Cluster analysis was performed using heatmap3 (Zhao, Guo et al., 2014) package in R. T-test was used to measure specific microbiome species abundance between conditions. Adjust p-value > 0.05 was used as significant threshold. Principle coordinate analysis was conducted in R. Confocal Images Statistical Analysis and Additional Software - Pearson’s Correlation Coefficient was acquired from BMM images using Huygens’s Deconvolution Scientific Volume Image Software (UNM Fluorescence Microscopy and Cell Imaging shared resource). Quantification figures were also made using Prism, while confocal image figures were constructed using Adobe Illustrator (version CC 2019). All other data were analyzed using one -way ANOVA or two-tailed unpaired Student’s t test (Prism).

## Online supplemental material

Material and methods for cell and tissue preparation, RT-qPCR, flow cytometry, histology, immunoblotting are found in supplemental material.

## Author contributions

SDM and SMG performed all analysis with the help from YG, ZERW, JRB, RRG, AJH, JGI and SBB. SDM, ZERW, YG, KCS and DLD contributed to microbiota sequencing and analysis. SMG, JRB and SBB participated in imaging and analysis. JGI, SBB and VD provided reagents and animals. SDM and SMG participated in writing the manuscript. EFC designed the study, analyzed data and wrote the paper. All authors approved the final version of the manuscript.

## Acknowledgments

Supported in part by the National Center for Research Resources and the National Center for Advancing Translational Sciences of the National Institutes of Health (NIH) through grant no. UL1TR001449 (E.F.C.) and in part by NIH grant P20GM121176 (E.F.C.), and the Bioinformatics Shared Resource at University of New Mexico Comprehensive Cancer Center with grant P30CA118100 (Y.G.). S.M.G. was supported in part by the Infectious Disease and Inflammation Program pre-doctoral T32 training grant, NIH/NIAID grant T32AI007538.

## Supplementary Information

### Cells and tissue preparation

Mesenteric lymph nodes (mLN) were collected and manually homogenized into single cell suspensions in RPMI prior to plating and 4-hour stimulation with PMA (50 ng/mL), Ionomycin (500 ng/mL), Brefeldin A/GolgiPlug (1:1000, BD Biosciences), and Monesin/GolgiStop, (1:1000, BD Biosciences) in preparation for intracellular cytokine staining and analysis by flow cytometry. Colon samples were cryopreserved in RNA-Later (Invitrogen), or cultured in RPMI for 4 hrs for cytokine analysis by ELISA. The histopathological evaluation was performed on the mouse colon sections using Hematoxylin and Eosin (H and E) stained slides after formalin fixation. Standard five-micron thick H and E sections obtained from paraffin embedded blocks fixed in 10% formalin were used for assessment. The large intestine of the mice was embedded in paraffin using the ‘swiss-roll’ technique. The generation of BMM were prepared as briefly described, marrow was collected from the femur and tibia, washed and differentiated in DMEM (Gibco), FBS (VWR), and murine L929-fibroblast supernatant containing CSF-1 for 10 days, after which cellular morphology was evaluated to confirm differentiation. BMM’s rested for 16 hours in media without CSF-1 prior to stimulation as specified for each experimental procedure below where relevant. All reagents used in cell isolation, culture, and stimulation are listed in Table S1. Flow cytometric protocol is described below, and all antibodies used for flow cytometric analysis are included in Table S7.

### RNA Isolation, Quantification, and RT-qPCR

RNA isolation was performed on cells or tissues cryopreserved in RNA-Later (Invitrogen) using the RNA Purelink Minikit (Invitrogen) according to the manufacturer’s protocols. RNA was quantified on Nanodrop2000 and all samples yielded a 260/280 of 2 ± 0.15. cDNA synthesis was performed using Oligo(dT) Primer and SSIV Reverse Transcriptase in the presence of Cloned Ribonuclease Inhibitor (all ThermoFisher). Reverse transcription reaction and RT-qPCR runs utilized Taqman MasterMix (ThermoFisher) on a viiA7 Thermal Cycler (ThermoFisher) using QuantStudio 7. RT-qPCR primers and reagents used are listed in Table S3.

### ClusPro web server

The ClusPro web server for protein-protein docking was utilized to assess the interaction between IL-12p (PDB ID: 1F45) and NBR1 (PDB IDs: 1WJ6, 2L8J, 2MGW, and 2MJ5).

https://cluspro.org/jobdetail.php?job=458409

https://cluspro.org/jobdetail.php?job=458405

https://cluspro.org/jobdetail.php?job=458404

https://cluspro.org/jobdetail.php?job=458403

### siRNA knockdown, Western blot, co-immunoprecipitation analysis and protein detection assays

Small interfering RNA knockdowns were achieved using Amaxa Mouse Macrophage NucleofectorTM Kit (Lonza). Nucleofection was performed on Nucleofector 2b Device (Lonza) per manufacturer’s protocols, using 20 μM reconstitutions of Non-Targeting (Dharmacon) and 5 μM reconstitutions of NBR1 (Dharmacon) siRNA. Following Nucleofection, cells were plated and rested for 16 hours prior to stimulation. All macrophages were stimulated with LPS (500 ng/mL), IFNy (10 ng/mL) for 6 hours, with one of two duplicates also receiving BafA1 for 5.5 hours. siRNA reagents are listed in Table S5. All ELISA were performed using Quantikine ELISA Kits (R&D) according to the manufacturer’s specifications. For all analyses, undiluted cell media was collected immediately prior to cell harvesting and placed on ice prior to freezing. ELISA quantification was performed on Synergy Neo2 (BioTek) and plate washes using Quantikine wash buffer were performed on Multiwash Advantage (Tricontinent). All reagents used were taken from the Quantikine kits listed in Table S6. For flow cytometry, cells were washed poststimulation with FACS Buffer [90% by volume 1x PBS, (Life Technologies), 10%FBS (VWR), and 0.05% 0.5 M EDTA, (Invitrogen)], and incubated for 20 min with antibody stain, followed by further FACS wash, and fixation in 1% PFA. To examine intracellular proteins, cells were permeabilize and incubated with intracellular cytokines and washed and resuspended FACS buffer. Sample analysis was performed on LSR Fortessa (BD Biosciences). Antibodies used for flow cytometric analysis are listed in Table S7. For immunoblots, after stimulation, cells were washed in 1x PBS and lysed in 20 uL of Pierce RIPA Lysis Buffer (ThermoFisher). All samples were boiled for 5 min at 65C in 1:1 Pierce RIPA Buffer: 2x Laemmli Buffer prior to gel electrophoresis at 90V (10% Mini-PROTEAN^®^ TGX™ Precast Protein Gels, Mini-PROTEAN^®^ Tetra Vertical Electrophoresis Cell for Mini Precast Gels, 2-gel, BioRad). Immunoblot transfer used the Trans-Blot^®^ Turbo™ Transfer System (Bio-Rad), and all blots were probed in 5% dry milk. All blots were imaged on Odyssey Clx (Licor Biosciences). For co-immunoprecipitated of IL-12p35 to NBR1 and IL-12p40 or NBR1 to IL-12p35 and IL-12p40, lysates were incubated with protein A/G magnetic beads with 2μg of anti-IL-12p35 or anti-NBR1 antibody or isotype control for 4 hours at 4°C. The bead-antibody complexes were washed with cold PBS followed by incubation with whole cell lysates overnight at 4°C. Following washes with lysis buffer and cold PBS, the bead-immune complexes were then resuspended in Laemmli’s sample buffer and boiled for 5 minutes. Samples were then prepared for immunoblot analysis. All antibodies used for Western Blot are listed in Table S8.

---

**Supplementary Figure 1.**
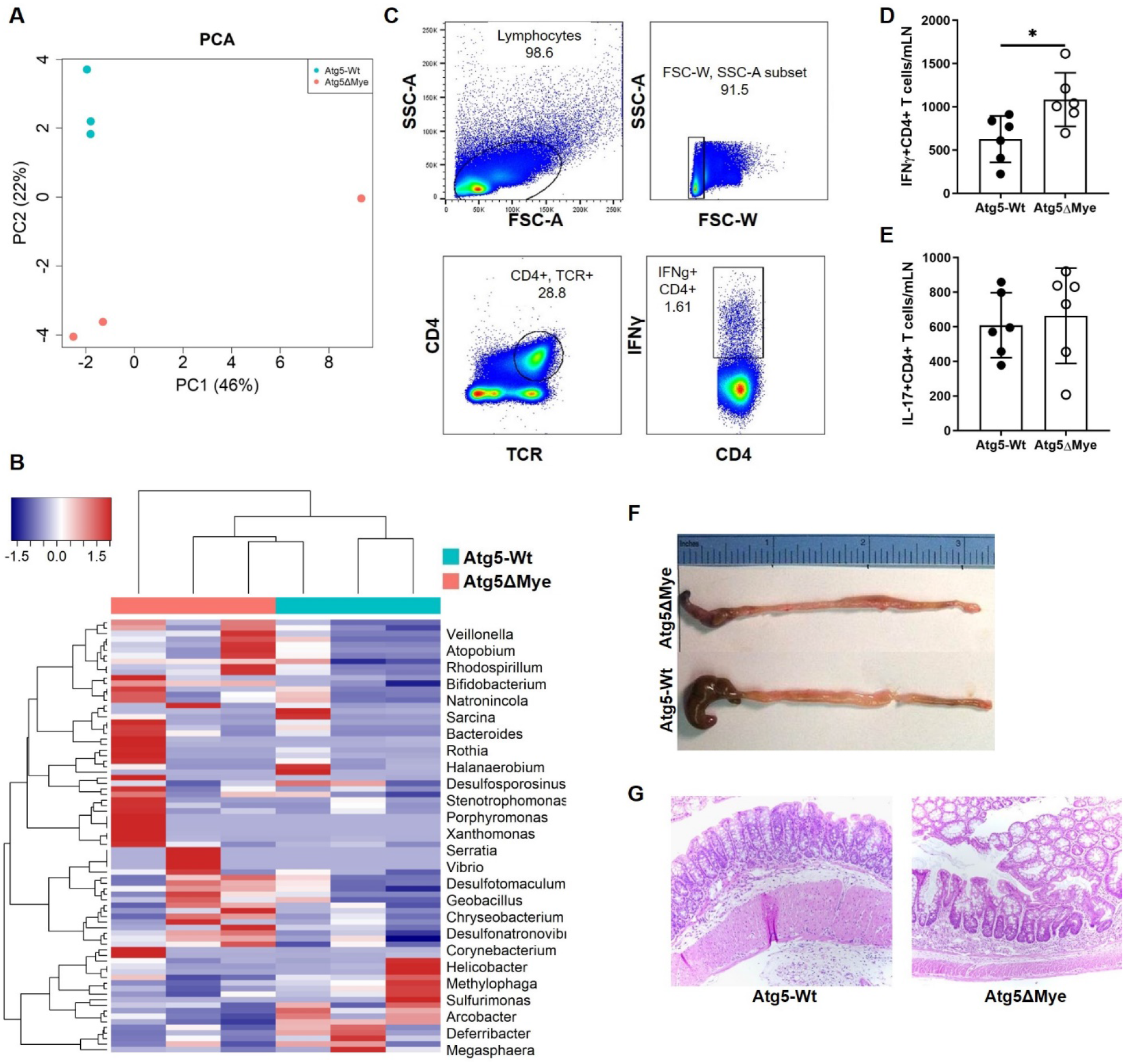
Steady State CD4^+^ T cell polarization and colitic response in Atg5cKO mice. **(A)** Principal coordinates analysis (PCoA) plot and **(B)** heatmap of microbiota composition in female mice (Genus level). **(C)** Flow cytometric gating of mLN CD4^+^ T cells for intracellular cytokine staining. Representative dot plots of intracellular IFNγ expression in mLN CD4^+^ T cells isolated from Atg5-Wt and Atg5ΔMye mice. **(D and E)** Graph of IFNγ^+^ and IL-17^+^ CD4^+^ T cells isolated from the mLN of Atg5-Wt and Atg5ΔMye mice. **(F)** Representative gross anatomy of the cecum and colon of colitic Atg5-Wt and Atg5ΔMye mice. **(G)** H&E staining of colons. Representative of two-three independent experiments, Graphs indicate mean (±SD). Student’s t test, P<0.05.

**Supplementary Figure 2.**
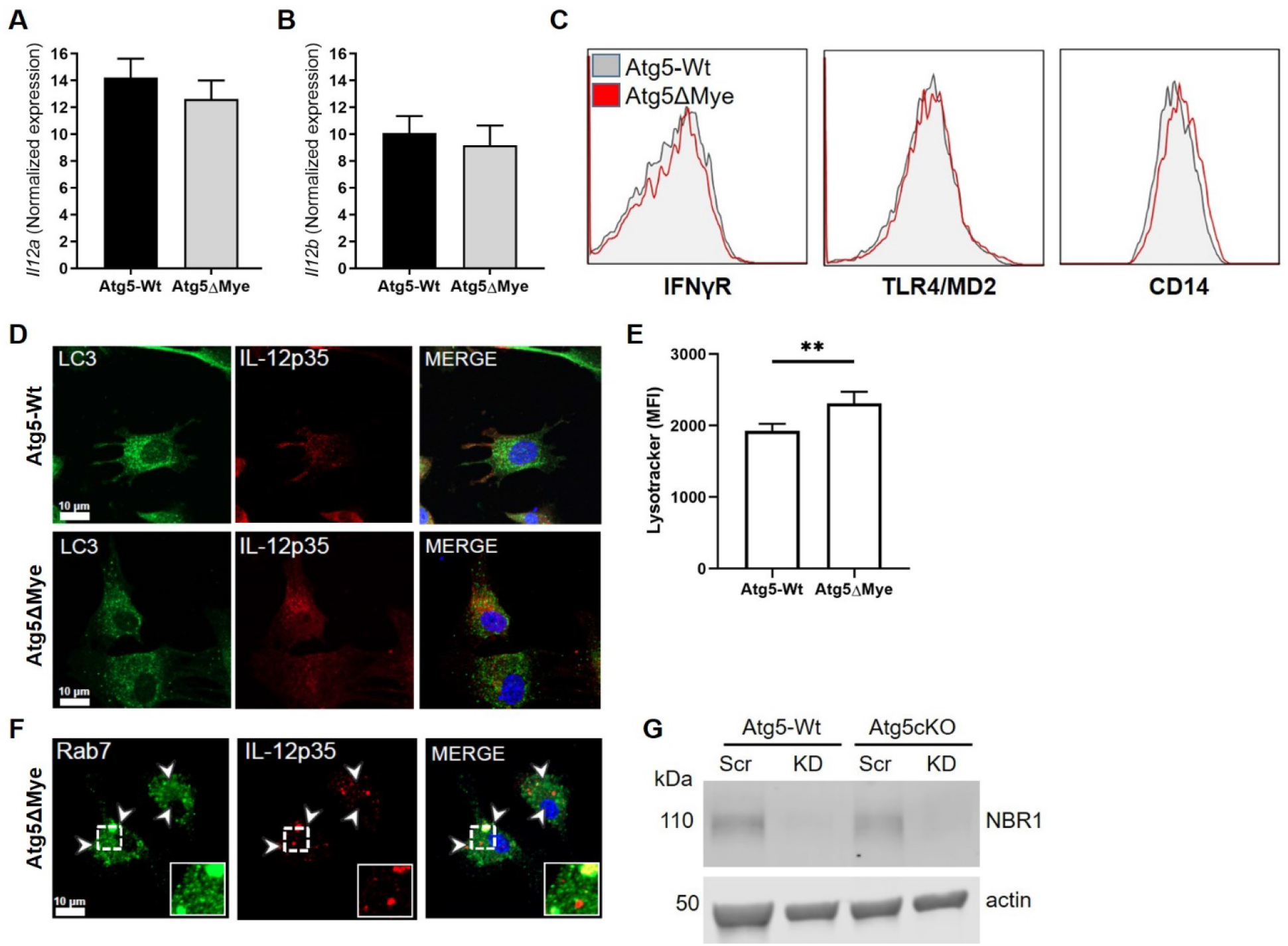
Regulation of IL-12 secretion in macrophages. **(A)** *Il12a* and **(B)** *Il12b* gene expression in unstimulated macrophages. **(C)** Representative histograms of cell surface expression on IFNγR (left), TLR4/MD2 (center), and CD14 (right) between unstimulated Atg5-Wt (gray) and Atg5ΔMye (red) macrophages. **(D)** Atg5-Wt and Atg5ΔMye macrophages were stained for LC3 (autophagosome marker) and IL-12p35 (subunit of cytokine IL12) after LPS, IFNγ and BafA1. Representative from 20 images from 5 slides. **(E)** Mean fluorescence intensity (MFI) of Lysotracker green in unstimulated Atg5-Wt and Atg5ΔMye macrophages. **(F)** Atg5ΔMye macrophages were stained for Rab7 (late endosome marker) and IL-12p35 (subunit of cytokine IL12) after LPS, IFNγ and BafA1. Immunoblot of IL-12p35 and actin expression in Atg5-Wt and Atg5ΔMye macrophages after stimulation with LPS and IFNγ or LPS, IFNγ and BafA1. Representative from 20 images from 3 slides. Arrows in images indicate puncta colocalizing with insets displaying enlargement of indicated region. **(G)** Immunoblot of NBR1 expression for scrambled (Scr) or knockdown (KD) siRNA targeted of NBR1 in Atg5-Wt and Atg5ΔMye macrophages after stimulation with LPS and IFNγ. Representative of two-three independent experiments, Graphs indicate mean (±SD).

**Supplementary Figure 3.**
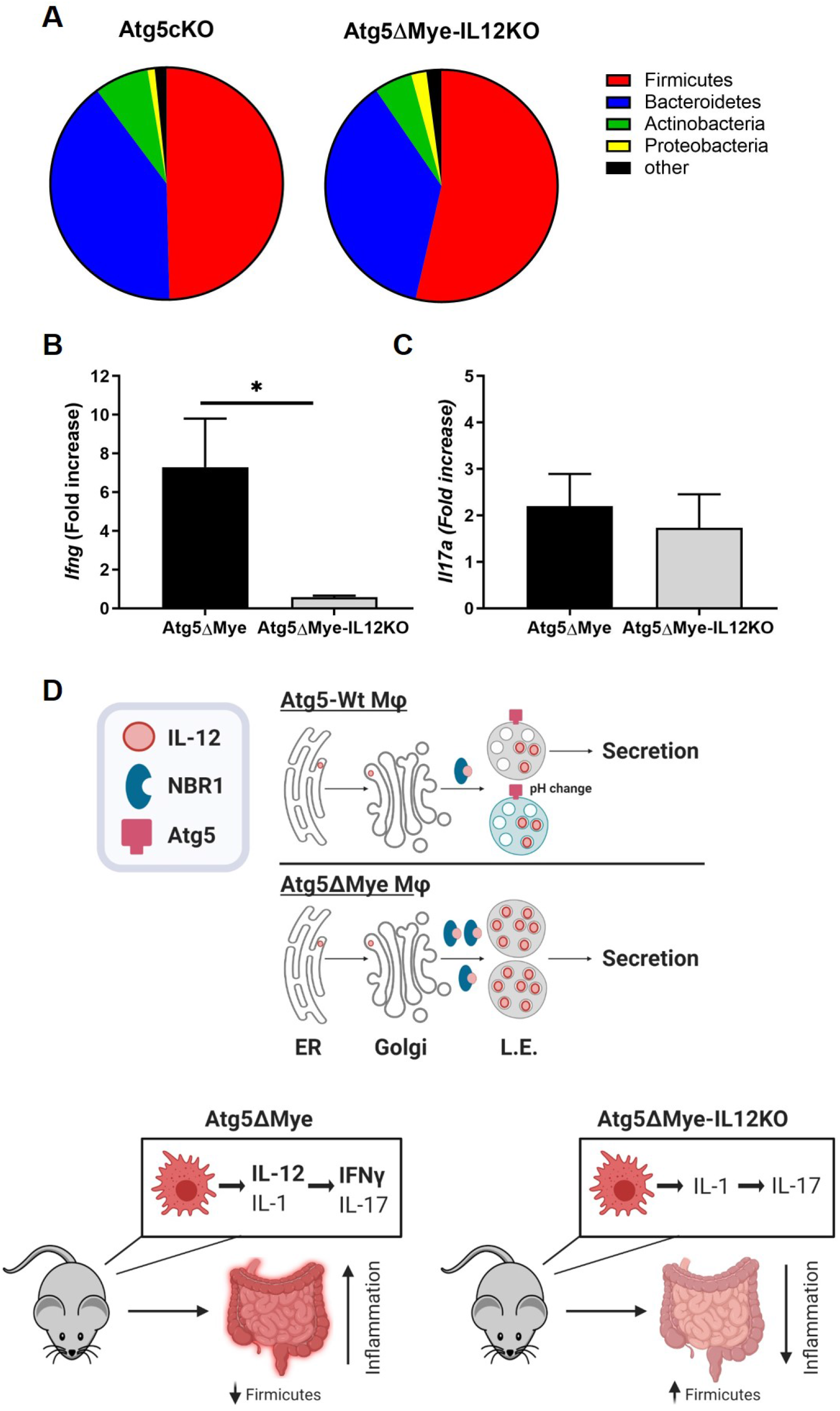
Comparison of Atg5ΔMye and Atg5ΔMye-IL12KO mice. Metagenomic sequencing and 16S profiling of DNA extracted from freshly collected stool of Atg5ΔMye and Atg5ΔMye-IL12KO mice (n=5 per group). **(A)** Pie chart showing the average proportion of Firmicutes, Bacteroidetes, Actinobacteria, Proteobacteria and all other phyla in 8-week old mice. **(B and C)** Colonic *Ifng* and *Il17a* gene expression after colitis induction. **(D)** Graphical abstract (BioRender). Representative of two independent experiments, Graphs indicate mean (±SD). * P<0.05, Two-tailed unpaired Student’s t tests.

